# Hierarchical Timescales in the Neocortex: Mathematical Mechanism and Biological Insights

**DOI:** 10.1101/2021.09.06.459048

**Authors:** Songting Li, Xiao-Jing Wang

## Abstract

A cardinal feature of the neocortex is the progressive increase of the spatial receptive fields along the cortical hierarchy. Recently, theoretical and experimental findings have shown that the temporal response windows also gradually enlarge, so that early sensory neural circuits operate on short-time scales whereas higher association areas are capable of integrating information over a long period of time. While an increased receptive field is accounted for by spatial summation of inputs from neurons in an upstream area, the emergence of timescale hierarchy cannot be readily explained, especially given the dense inter-areal cortical connectivity known in modern connectome. To uncover the required neurobiological properties, we carried out a rigorous analysis of an anatomically-based large-scale cortex model of macaque monkeys. Using a perturbation method, we show that the segregation of disparate timescales is defined in terms of the localization of eigenvectors of the connectivity matrix, which depends on three circuit properties: (1) a macroscopic gradient of synaptic excitation, (2) distinct electrophysiological properties between excitatory and inhibitory neuronal populations, and (3) a detailed balance between long-range excitatory inputs and local inhibitory inputs for each area-to-area pathway. Our work thus provides a quantitative understanding of the mechanism underlying the emergence of timescale hierarchy in large-scale primate cortical networks.

**Significance Statement:** In the neocortex, while early sensory areas encode and process external inputs rapidly, higher association areas are endowed with slow dynamics suitable for accumulating information over time. Such a hierarchy of temporal response windows along the cortical hierarchy naturally emerges in a model of multi-areal primate cortex. This finding raises the question of why diverse temporal modes are not mixed in roughly the same way across the whole cortex, despite high connection density and an abundance of feedback loops. We investigate this question by mathematically analyzing the anatomically-based network model of macaque cortex, and show that three general principles of synaptic excitation and inhibition are crucial for timescale segregation in a hierarchy, a functionally important characteristic of the cortex.

---

THE brain is organized with a delicate structure to integrate and process both spatial and temporal information received from the external world. For spatial information processing, neurons along cortical visual pathways possess increasingly large spatial receptive fields, and its underlying mechanism has been understood as neurons in higher level visual areas receive input from many neurons with smaller receptive fields in lower level visual areas, thereby aggregating information across space (Hubel, 1995). More recently, a computational model (Chaudhuri et al., 2015) revealed that the timescale over which neural integration occurs also gradually increases from area to area along the cortical hierarchy. The model was based on the anatomically measured directed- and weighted-inter-areal connectivity of the macaque cortex (Markov et al., 2014a) and incorporated heterogeneity of synaptic excitation calibrated by spine count per pyramidal neuron (Elston, 2007). It has been observed that the decay times increased progressively along the cortical hierarchy when signals propagate in the network, and the temporal hierarchy could change dynamically in response to different types of sensory inputs (e.g., different hierarchy of timescales for somatosensory input versus visual input) (Chaudhuri et al., 2015). By manipulating parameters of the model, simulation results further demonstrate that both within and between regions of anatomical properties could affect the hierarchy of timescales in neuronal population activity (Chaudhuri et al., 2015). A hierarchy of temporal receptive windows is functionally desirable, so that the circuit dynamics operate on short time scales in early sensory areas to encode and process rapidly changing external stimuli, whereas parietal and frontal areas can accumulate information over a relatively long period of time window in decision-making and other cognitive processes (Gold and Shadlen, 2007; Wang, 2008).

Despite the accumulating evidence in support of timescale hierarchy across cortical areas in mice (Runyan et al., 2017; Siegle et al., 2021), monkeys (Ogawa and Komatsu, 2010; Murray et al., 2014; Cavanagh et al., 2016; Fascianelli et al., 2019; Spitmaan et al., 2020; Maisson et al., 2021; Manea et al., 2021), and humans (Hasson et al., 2008; Honey et al., 2012; Lerner et al., 2011; Stephens et al., 2013; Yeshurun et al., 2017; Raut et al., 2020; Shafiei et al., 2020; Gao et al., 2020), its underlying mechanism remains unclear. In particular, since inter-areal connections are dense, with roughly 65% of all possible connections present in the macaque cortex (Markov et al., 2014a) and even higher connection density in the mouse cortex (Gamanut et al., 2018), what circuit properties are required to ensure that dynamical modes with disparate time constants are spatially localized? How intra-areal anatomical properties determine the intrinsic timescale of each area, and how these intrinsic timescales remain to be segregated rather than mixed up in the presence of dense inter-areal connections? In this work, we addressed these questions by a mathematical analysis of the model (Chaudhuri et al., 2015). Using a perturbation method, we identified key required conditions, in particular a detailed excitation-inhibition balance for long-distance inter-areal connections which is experimentally testable.

## The multi-areal model and hierarchical timescales phenomenon

We first review the mathematical form of the multi-areal model of the macaque cortex, and the hierarchical timescales phenomenon captured by this model (Chaudhuri et al., 2015). The macaque cortical network model contains a subnet of 29 areas widely distributed from sensory to association areas in the macaque cortex, and each area includes both excitatory and inhibitory neuronal populations. The neuronal population dynamics in the *i*th area is described as

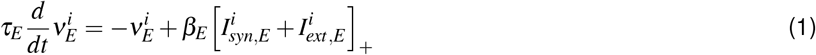

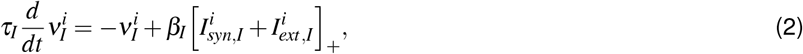

where 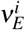 and 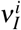 are the firing rate of the excitatory and inhibitory populations in the *i*th area, respectively, *τ_E_* and *τ_I_* are their time constants, respectively, and *β_E_* and *β_I_* are the slope of the f-I curve for the excitatory and inhibitory populations, respectively. The f-I curve takes the form of a rectified linear function with [*I*]_+_ = *max*(*I*, 0). In addition, 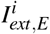 and 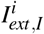 are the external currents, and 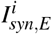 and 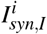 are the synaptic currents that follow

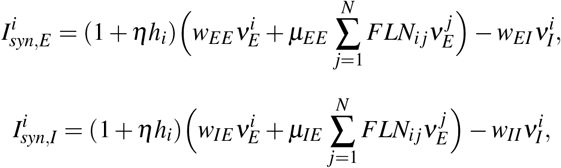

where *w_pq_, p, q* ∈ {*E, I*} is the local coupling strength from the *q* population to the *p* population within each area. *FLN_ij_* is the fraction of labeled neurons (FLN) from area *j* to area *i* reflecting the strengths of long-range input (Markov et al., 2014a), and *μ_EE_* and *μ_IE_* are scaling parameters that control the strengths of long-range input to the excitatory and inhibitory populations, respectively. Both local and long-range excitatory inputs to an area are scaled by its position in the hierarchy quantified by *h_i_* (a value normalized between 0 and 1), based on the observation that the hierarchical position of an area highly correlates with the number of spines on pyramidal neurons in that area (Elston, 2007; Chaudhuri et al., 2015). A constant *η* maps the hierarchy *h_i_* into excitatory connection strengths. Note that both local and long-range projections are scaled by hierarchy, rather than just local projections, following the observation that the proportion of local to long-range connections is approximately conserved across areas (Markov et al., 2011). The values of all the model parameters are specified in Sec. *Materials and Methods*.

By simulating the model, it has been observed in Ref. (Chaudhuri et al., 2015) that the decay time of neuronal response in each area increases progressively along the visual cortical hierarchy when an pulse input is given to area V1, as shown here in Fig. 1A. Early visual areas show fast and transient responses while prefrontal areas show slower responses and longer integration times with traces lasting for several seconds after the stimulation. In addition, white-noise input to V1 is also integrated with a hierarchy of timescales by computing the autocorrelation of neuronal activity at each area (Chaudhuri et al., 2015). As shown in Fig. 1B, the activity of early sensory areas shows rapid decay of autocorrelation with time lag while that of association areas shows slow decay. In Fig. 1C, by fitting single or double exponentials to the decay of the autocorrelation curves (Chaudhuri et al., 2015), the dominant timescale of each area tends to increase along the hierarchy approximately, thus a hierarchy of widely disparate timescales emerges from this model. It is worth noting, however, that the timescale does not change monotonically with the anatomically defined hierarchy (x-axis), the precise pattern is sculpted by the measured inter-areal wiring properties.

**Figure 1:**
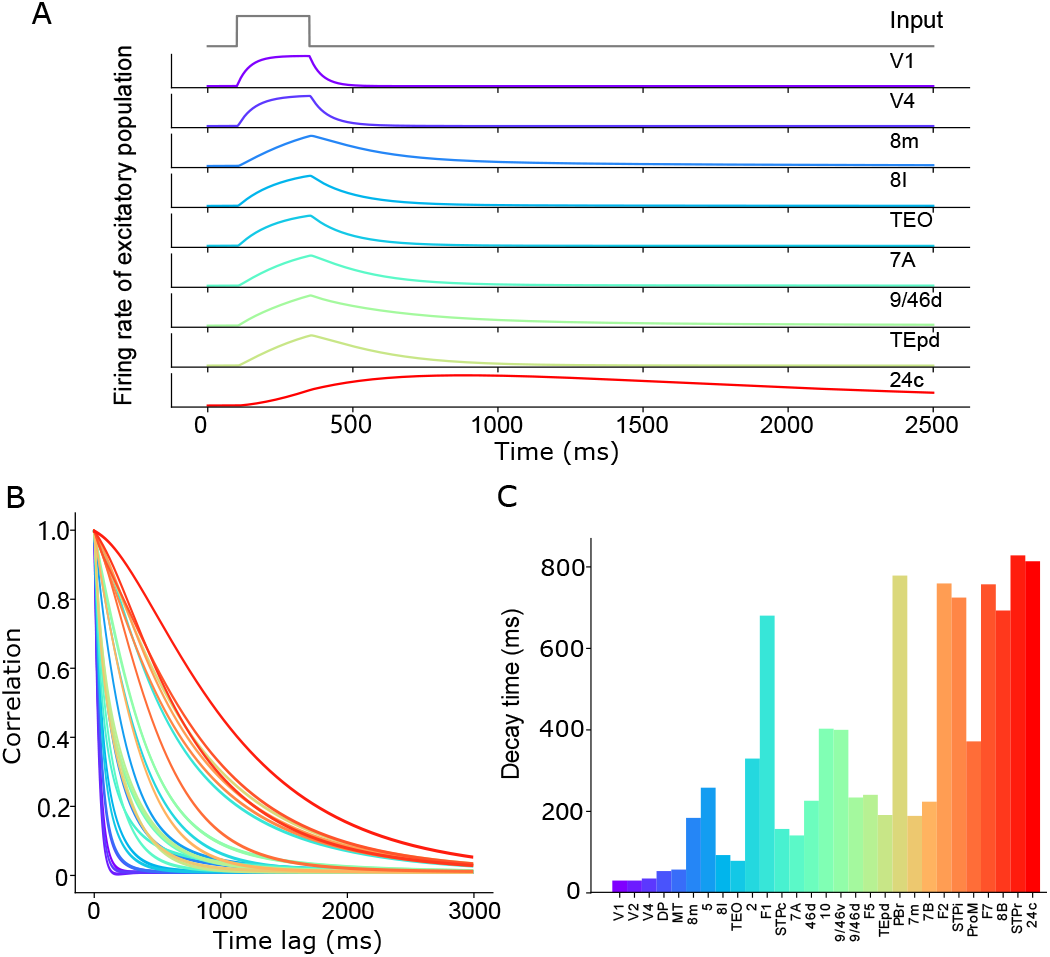
The hierarchical timescales phenomenon simulated in the macaque multi-areal model. (A), a pulse of input to area V1 is propagated along the hierarchy, displaying increasing decay times as it proceeds. (B), autocorrelation of area activity in response to white-noise input to V1. (C), the dominant time constants in all areas, extracted by fitting single or double exponentials to the autocorrelation curves (Chaudhuri et al., 2015). In (A)-(C), areas are arranged and colored by position in the anatomical hierarchy.

Note that, although the multi-areal model (Eqs. 1–2) is nonlinear by taking into account a rectified linear f-I curve, the stimuli in our simulations drive all neuronal population activities above the firing threshold with positive input currents to all areas. Therefore, the stimuli essentially drive the network dynamics into the linear regime. Before we perform mathematical analysis to understand the mechanism underlying the emergence of hierarchical timescales in the simulations, to simplify the notation, we rewrite the network dynamics Eqs. 1–2 in the linear regime in the following form

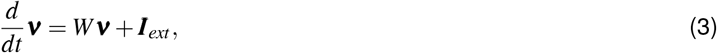

where

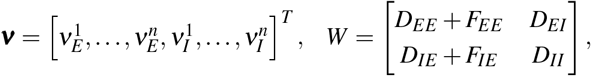

with *n* = 29, and *D_EE_*, *D_EI_*, *D_IE_*, *D_II_* being four diagonal matrices whose *i*th element on their diagonal line are

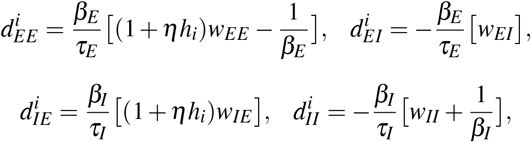

respectively, and matrices *F_EE_* and *F_IE_* being two non-diagonal matrices whose *i*th-row-*j*th-column element is

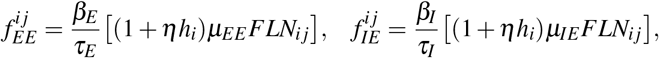

respectively. Note that matrices *D_EE_*, *D_EI_*, *D_IE_*, *D_II_* reflect local intra-areal interactions, while matrices *F_EE_* and *F_IE_* reflect long-range inter-areal interactions. In addition, elements in *D_EE_*, *D_IE_*, *F_EE_*, and *F_IE_* depend on area hierarchy *h_i_* while elements in *D_EI_* and *D_II_* are constant. Finally, the external input vector is

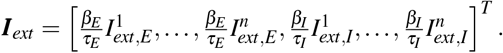

Denoting the eigenvalues and eigenvectors of the connectivity matrix *W* as *λ_i_* and ***V**_i_* (*i* = 1,2, …, 2*n*), respectively, i.e., *W**V**_i_* = *λ_i_**V**_i_*, the analytical solution of Eq. 3 can be obtained as

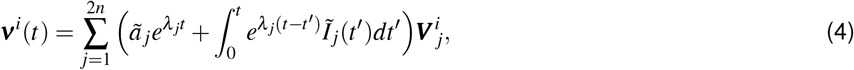

where ***ν**^i^* and 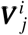 are the *i*th element in ***v*** and ***V**_j_*, respectively, 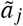 and 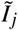 are the coefficients for the initial condition and the external input, respectively, represented in the coordinate system of the eigenvectors {***V**_j_*}. Note that, from Eq. 4, each area integrates input current with the same set of time constants {*τ_i_*} determined by the real part of the eigenvalues, i.e., *τ_i_* = −1/*Re*{*λ_i_*}. Therefore, the characteristic timescale of each area across the network is expected to be similar in general case. To obtain distinct timescales at each area, it requires (1) the localization of eigenvectors ***V**_j_*, i.e., most of the elements in ***V**_j_* are close to zero, (2) the orthogonality of all pairs of eigenvectors, i.e., the non-zero elements are nearly non-overlap for different ***V**_j_*.

By computing the eigenvalues and eigenvectors of matrix *W*, as shown in Fig. 2A, the timescale pool {*τ_i_*} derived from eigenvalues can be classified into two groups, one group shows quite fast timescale about 2 milliseconds, and the other group includes relatively slow timescales ranging from tens to hundreds of milliseconds. In addition, we are particularly interested in the excitatory population because the majority of neurons in the cortex are excitatory neurons. We observe that the magnitude of the eigenvectors corresponding to the fast timescale is nearly zero for the excitatory population at each area, while that corresponding to the slow timescale is weakly localized and weakly orthogonal, i.e., each eigenvector has a few nonzero elements that almost do not overlap with other eigenvectors’ nonzero elements. According to Eq. 4, the pattern of eigenvectors gives rise to the disparate timescales for the excitatory neuronal population at each cortical area. We next perform mathematical analysis to investigate the sufficient conditions for (1) vanishing magnitude of the excitatory component of fast-eigenmode eigenvectors, and (2) weak localization and orthogonality of the excitatory component of slow-eigenmode eigenvectors in this network system.

**Figure 2:**
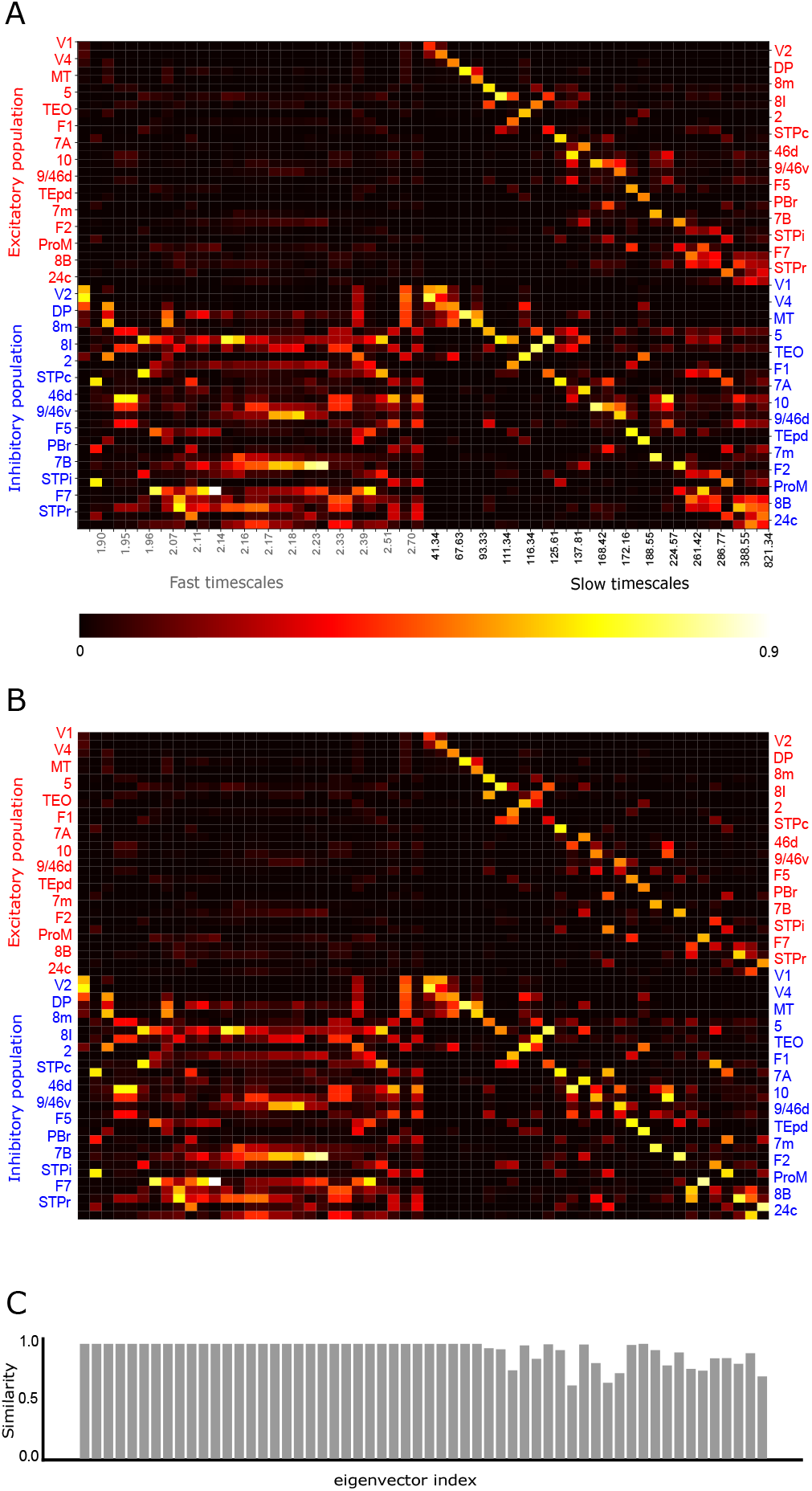
Eigenvectors of the network connectivity matrix and their approximations from the perturbation analysis. (A), eigenvectors of the network connectivity matrix *W*. Each column shows the amplitude of an eigenvector at the 29 areas, with corresponding timescale labeled below. (B), eigenvectors of *W* calculated from the first-order perturbation analysis. (C), similarity measure defined as the inner product of the corresponding eigenvectors in (A) and (B).

## Perturbation analysis of the model

We note that the parameters of the model (specified in Sec. *Materials and Methods*) gives

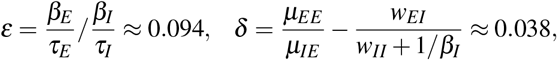

which can be viewed as two small parameters in order to allow us to perform perturbation analysis below.

We first study the network in the absence of the long-range interactions among areas. In this scenario, we study the 2 by 2 block matrix 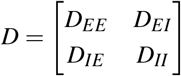 in which each block is a diagonal matrix defined above. By viewing 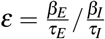 as a small parameter, we have

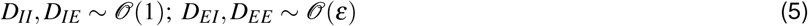

from their definitions. Accordingly, we can prove the following proposition:

### Proposition 1

*If* 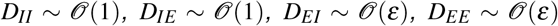, *then D has n eigenvalues being* 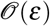, *and n eigenvalues being* 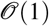.

**Proof**: It is straightforward to prove that Matrix *D* can be diagonalized by matrix *P*, i.e.,

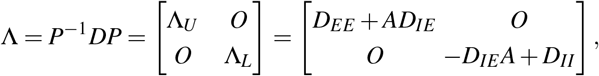

where

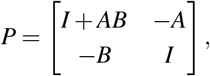

*I* is the identity matrix, and diagonal matrix *A* satisfies −(*D_EE_* + *AD_IE_*)*A* + *D_EI_* + *AD_II_* = 0, diagonal matrix *B* satisfies *D_IE_* + *B*(*D_EE_* + *AD_IE_*) + (*D_IE_A* – *D_II_*)*B* = 0.

We solve the equation of *A* and choose one of the two solutions of *A* as

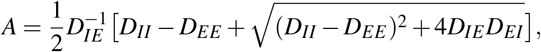

where the square root of a diagonal matrix is defined as taking the square root of its elements. Due to the fact that 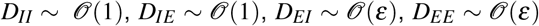, we have 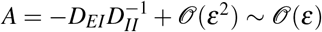, and 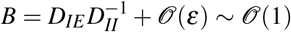. Accordingly, we have

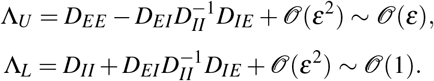

From Proposition 1, the eigenvalues of *D* have two separated scales belonging to Λ*_U_* and Λ_*L*_, respectively. As the timescales of the network system is given by *τ_i_* = −1/*Re*{*λ_i_*} (*λ_i_* is the *i*th diagonal element in matrix Λ), the separation of scales for eigenvalues in Λ*_U_* and Λ*_L_* explains that the intrinsic timescale pool can be classified into two group with a separation of scales, which is mainly determined by the distinct electrophysiological properties between excitatory and inhibitory neuronal populations within each area described by *ε*. In addition, from the analysis, the eigenvalues in Λ*_L_* with large magnitude (fast timescale) are less sensitive to the hierarchy level because Λ*_L_* ≈ *D_II_*, and the elements in *D_II_* does not depend on *h_i_*. Therefore, the gradient of *h_i_* across areas barely affects fast timescale pool. In contrast, the eigenvalues in Λ_*U*_ with small magnitude (slow timescale) are more sensitive to the hierarchy level because 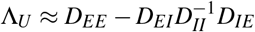, and both the elements in *D_EE_* and *D_IE_* depend on *h_i_*. Therefore, the gradient of *h_i_* across areas increases the range of slow timescale pool. Further, the slow timescales of each area in this disconnected network are segregated and follows the hierarchical order *h_i_* as the corresponding eigenvectors are perfectly localized and orthogonal to each other.

Now we consider the multi-areal network in the presence of long-range interactions. Adding long-range connectivity to local connectivity matrix *D* changes the eigenvalues and eigenvectors of matrix *D*, which can be analyzed in the following.

By multiplying *P* and *P*^−1^ (given in the proof of Proposition 1) on both sides of *W*, we have

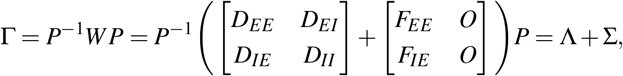

where

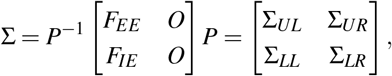

with 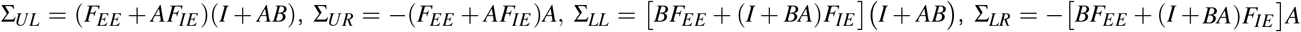, and with Λ, *A*, and *B* defined in the proof of Proposition 1.

Denoting one of the eigenvectors of matrix Γ as [***u*, *v***]*^T^*, and the corresponding eigenvalue as *λ*, we have

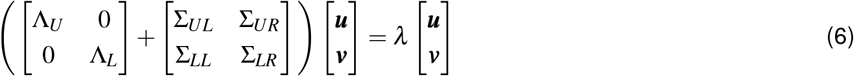

According to the definition of 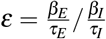, and 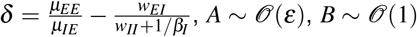, and 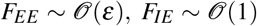, we have 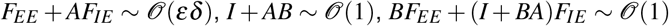, and accordingly,

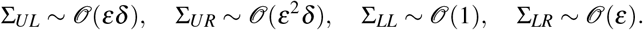

As 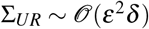 is a higher order term compared with Λ*_U_*, Λ*_L_*, Σ*_UL_*, Σ*_LL_*, and Σ*_LR_*, it can be dropped out in Eq. 6 and the error of eigenvalue and eigenvector is at most 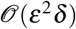 (see Proposition 4 in *Appendix* for a detailed proof). Accordingly, we can obtain two equations from Eq. 6 in the vector form,

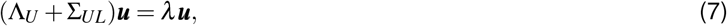

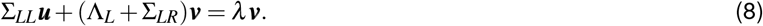

To describe the eigenvector property of Eqs. 7–8, we first introduce the definitions of weak localization and weak orthogonality as follows.

### Definition 1

*A vector **u***(*δ*) *is weakly localized if it can be represented as* 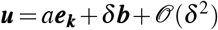 *for some k, where* 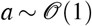 *is a constant number*, 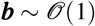 *is a constant vector, δ is a small parameter, and **e**_k_ represents the natural basis with only the kth element being 1 and others being zero, i.e., **e**_k_* = [0, …, 1(*kth*), …, 0].

### Definition 2

*Two vectors **u***(*δ*) *and **v***(*δ*) *are weakly orthogonal to each other if their inner product* 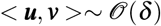, *where δ is a small parameter*.

With the concept of weak localization and weak orthogonality defined above, we introduce the following proposition that describes the property of ***u*** in the system of Eqs. 7–8.

### Proposition 2

*In the system described by Eqs. 7–8, if all matrices are analytic with respect to ε and δ, and if* 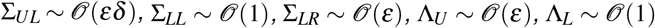, *and if* Λ*_U_ has n simple eigenvalues, then*

1. *there exists n eigenvectors* [***u, v***]*^T^ in which u =* 0, *with* 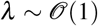 *correspondingly*.
2. *there exists n eigenvectors* [***u, v***]*^T^ in which **u** is weakly localized and weakly orthogonal to each other, with* 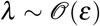 *correspondingly*.

**Proof:**

1. It is noted that ***u*** = 0 is a trivial solution of Eq. 7. By defining 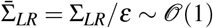, Eq. 8 becomes

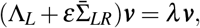

in which ***v*** is the eigenvector of matrix 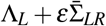. By viewing 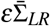 as a perturbation matrix to Λ*_L_*, then the leading order of *λ* shall be the same as that of *n* elements in the diagonal line of Λ_*L*_, which takes the order of 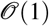.
2. In Eq. 7, if ***u*** ≠ 0, by defining 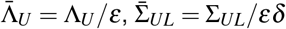, and 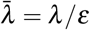, we have

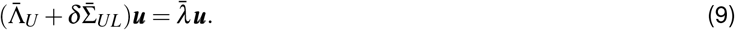 Therefore, ***u*** is also the eigenvector of matrix 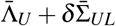, and 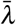 is the corresponding eigenvalue. As Λ*_U_* has *n* simple eigenvalues, so does 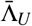, then ***u*** and 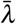 are analytic with respect to the perturbation parameter *δ* (Kato, 1966), i.e., 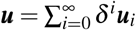, and 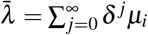 for *δ* near zero. Therefore, to the leading order, we have

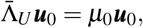

in which *μ*_0_ is the eigenvalue of the diagonal matrix 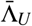, and ***u***_0_ is the corresponding eigenvector. Accordingly, ***u***_0_ ∈ {***e**_k_*}, thereafter

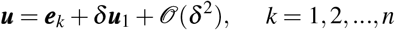

where ***e**_k_* represents the *k*th natural basis, and the leading order of 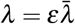 is 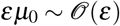. It’s straightforward to verify that ***u*** are weakly localized and weakly orthogonal to each other.

If we denote the unit-length eigenvector of the connectivity matrix *W* as [***r**_E_, **r**_I_*]*^T^*, and denote the corresponding eigenvalue as *λ* (the same as that of matrix Γ after the similarity transform), then from Propositions 1 and 2, we have

### Proposition 3

*Under the same conditions in Propositions 1-2, the unit-length eigenvector* [***r**_E_, **r**_I_*]*^T^ of the connectivity matrix W has the following properties*,

1. *for eigenvalue* 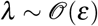, *the corresponding **r**_E_ is weakly localized and weakly orthogonal to each other*.
2. *for eigenvalue* 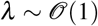, *the corresponding* 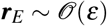.

**Proof:** We first consider [***r**_E_, **r**_I_*]*^T^* with non-unit length. According to the similarity transform, we have the following linear relation between [***r**_E_, **r**_I_*]*^T^* and [***u***, ***v***]^*T*^,

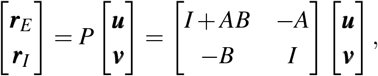

i.e., ***r**_E_* = (*I*+*AB*)***u*** – *A**v***, and ***r**_I_* = – *B**u*** + ***v***. From Proposition 1, we have 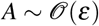, and 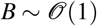.

1. From Proposition 2, we have 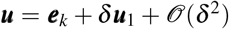 for 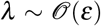 (*k* = 1,2,…, *n*). Accordingly, ***v*** can be solved as 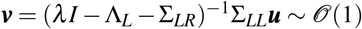, which gives 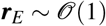, and 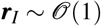. Therefore, the length of [***r**_E_, **r**_I_*]*^T^* denoted by *c* is order 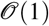. By normalizing the length of [***r**_E_, **r**_I_*]*^T^* to be unity, we have 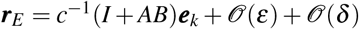 being weakly localized and weakly orthogonal to each other.
2. From Proposition 2, we have 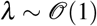. Accordingly, ***v*** can be solved as the eigenvector of matrix (Λ*_L_* + Σ*_LR_*) with unit-length. Therefore, [***r**_E_, **r**_I_*]*^T^* = [−*A**v, v***]^*T*^, and the length of [***r**_E_, **r**_I_*]*^T^* denoted by *c* is order 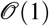. By normalizing the length of [***r**_E_, **r**_I_*]*^T^* to be unity, we have 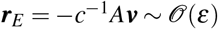.

Note that, Propositions 2-3 hold for sufficiently small *ε* and *δ* near zero. However, the convergence radius of the power series of ***u*** in Proposition 2 is not specified yet. Although difficult to calculate the convergence radius, we can compute the analytical expression of ***u***_1_ in the power series 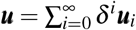 in Proposition 2 to obtain the first-order perturbation solution of ***u*** and thereby ***r**_E_*, which could help us gain insight about when weak localization and orthogonality of ***u*** and ***r**_E_* will break down approximately.

To the order of *δ* in Eq. 9, we have

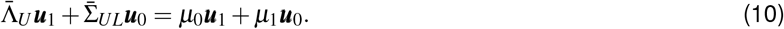

Without loss of generality, we assume ***u***_0_ = ***e**_k_*, and accordingly, 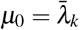 is the *k*th element in the diagonal line of matrix 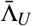. In addition, we normalize ***u*** to make the *k*th element in ***u*** denoted by ***u**^k^* to be unity, correspondingly 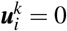 for *i* ≥ 1. By left multiplying 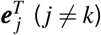 to Eq. 10, we have

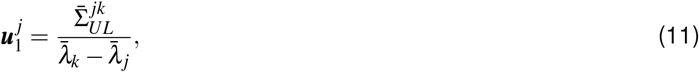

where 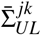 is the element in the *j*th row and *k*th column in matrix 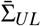. To make the first-order perturbation solution valid, we expect that 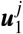 is small compared with *δ*^−1^, otherwise the separation of orders will no longer hold in the power series (i.e., first-order term *δ**u***_1_ becomes larger than the zeroth-order term ***e**_k_*). In such a case, the spectral gap 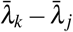 shall be large enough compared with elements in 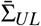. The spectral gap of matrix 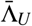 attributes to the gradient of excitation across areas, or simply *h_i_*. In Fig. 2B-2C, we show that the eigenvector [***r**_E_, **r**_I_*]*^T^*, which is solved using the perturbation theory in Proportions 2-3 to the first-order accuracy (Eq. 11), well agrees with the original eigenvector in most cases. However, some eigenvectors show less similarity to the original eigenvectors when the first-order perturbation theory breaks down for the reason discussed above.

## Biological interpretations of the three requirements

From the above analysis, three conditions are required to obtain weakly localized and orthogonal eigenvectors in order to maintain the hierarchy of timescales: (1) small *ε*, (2) small *δ*, (3) the gradient of *h_i_* across areas. We briefly summarize the important roles of the three conditions on proving eigenvector localization and orthogonality in the perturbation analysis illustrated in Fig. 3. As shown in Fig. 3A-3B, to remove intra-areal interactions between the excitatory and inhibitory populations within each area, we first change the coordinate system from (***r**_E_, **r**_I_*) to (***u, v***) with a transform matrix *P* given in Proposition 1. In the new coordinate system, there is no local interaction between the dynamical variables ***u*** and ***v***. Furthermore, considering a directed long-range projection from area *j* to area *i* in Fig. 3C, it has been shown that small *δ* leads to weak interaction from *u^j^* to *u^i^*, and small *ε* additionally leads to even weaker interaction from *v^j^* to *u^i^* that can be removed with an ignorable error (see Proposition 4 in Appendix). And a gradient of *h_i_* leads to a nonzero spectral gap between area *i* and area *j*. All the three conditions result in the weak localization and orthogonality of the ***u*** component in the (***u, v***) coordinate system, as proved by Proposition 2 and Eq. 11. Finally, as shown in Fig. 3D, small *ε* gives rise to the fact that the leading order of ***u*** and ***r**_E_* are the same, thus ***r**_E_* is also weakly localized and orthogonalized similar to ***u***, as proved by Proposition 3.

**Figure 3:**
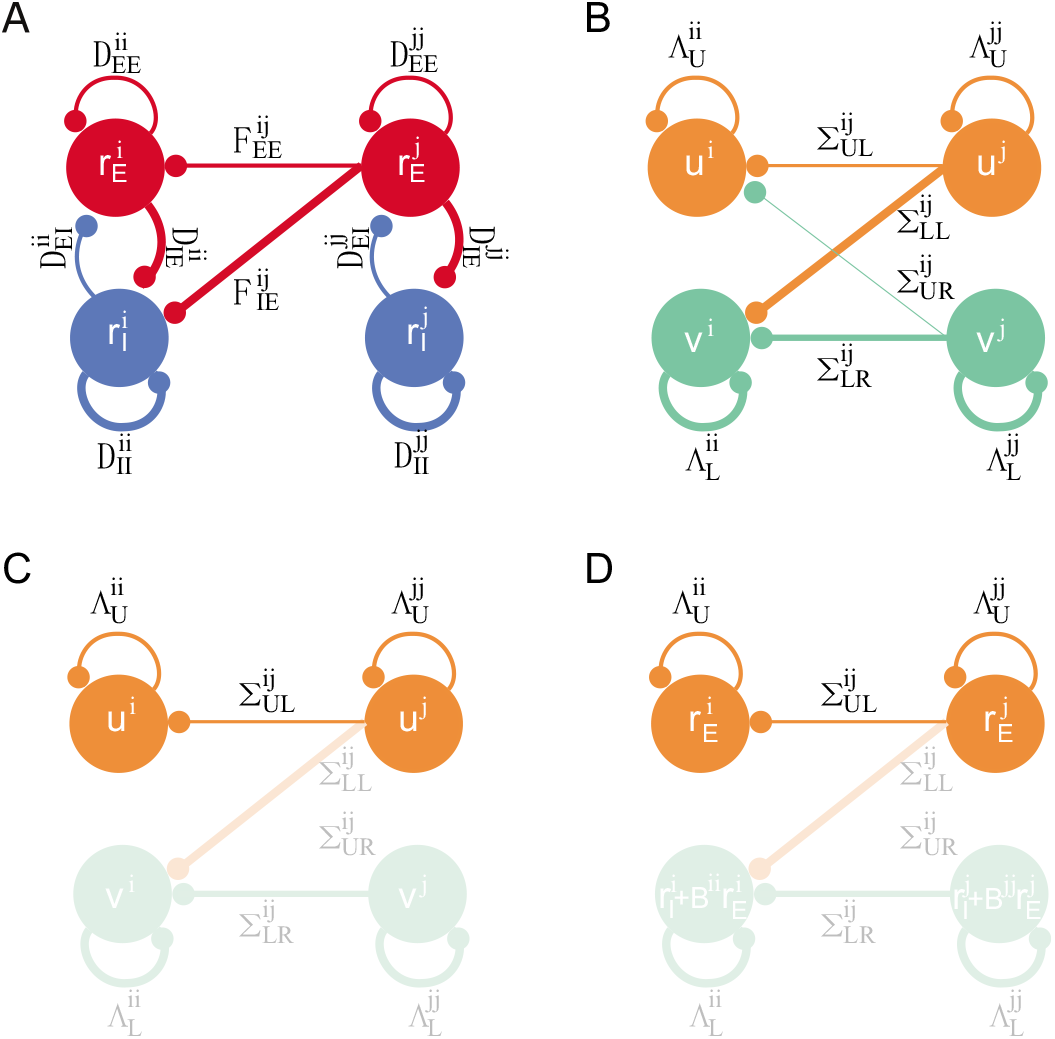
Schematics for the proof of weakly localized and orthogonal eigenvectors of the connectivity matrix *W*. (A), directed interaction from area *j* to area *i* in the original model (Eqs. 1–2). (B), one-way interaction from area *j* to area *i* after changing the coordinate system from (***r**_E_, **r**_I_*) to (***u, v***). (C), small *δ* leads to weak interaction from *u^j^* to *u^i^*, small *ε* additionally leads to even weaker interaction from *v^j^* to *u^i^* that is ignorable (proved by Proposition 4 in Appendix), and a gradient of *h_i_* leads to a nonzero spectral gap between area *i* and area *j*. Accordingly, they together lead to the weak localization and orthogonality of the ***u*** component in the (***u, v***) coordinate system (proved by Proposition 2 and Eq. 11). (D), one-way interaction from area *j* to area *i* after changing the coordinate system from (***u, v***) back to (***r**_E_, **r**_I_*). To the leading order, one has 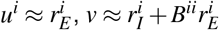. In this step, small *ε* ensures that the leading order of ***u*** and ***r**_E_* are identical, so are their localization and orthogonality properties (proved by Proposition 3). In (A)-(D), the width of lines codes the interaction strength, and light colored lines and nodes are not important in the proofs.

We next discuss the biological interpretation of the three conditions. First, according to the definition of 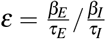, small *ε* indicates that the electrophysiological properties of excitatory and inhibitory neurons are different, in particular, their membrane time constant and the slope of the gain function. The substantial difference of electrophysiological properties between the excitatory and inhibitory neurons has been supported by experimental evidence, i.e., inhibitory neurons have larger slope of the gain function and smaller membrane time constant (Ahmed et al., 1998; Nowak et al., 2003; Povysheva et al., 2008; Zaitsev et al., 2012).

Second, small 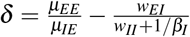 indicates the balanced condition between the inter-areal excitatory and intra-areal inhibitory inputs. When the presynaptic excitatory input from the *j*th area is increased by 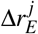, its influence on the excitatory population activity in the *i*th area in the steady state can be calculated in a straightforward way as 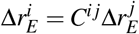, in which 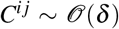. This indicates that the signal from the *j*th area have a small influence on the activity of the excitatory population in the *i*th area, because the global long-range excitatory input is balanced with and canceled by the local inhibitory synaptic input, leading to small net inputs in each signal pathway, as shown in Fig. 4. This condition corresponds to a detailed balance of excitation and inhibition that may benefit signal control and gating, as proposed in previous studies (Vogels and Abbott, 2009). The importance of excitation-inhibition balance on timescale hierarchy is supported by a recent study showing that the imbalance of excitation and inhibition could have a substantial effect on the change of intrinsic timescales across brain areas, which is a manifestation of psychosis such as hallucination and delusion (Wengler et al., 2020).

**Figure 4:**
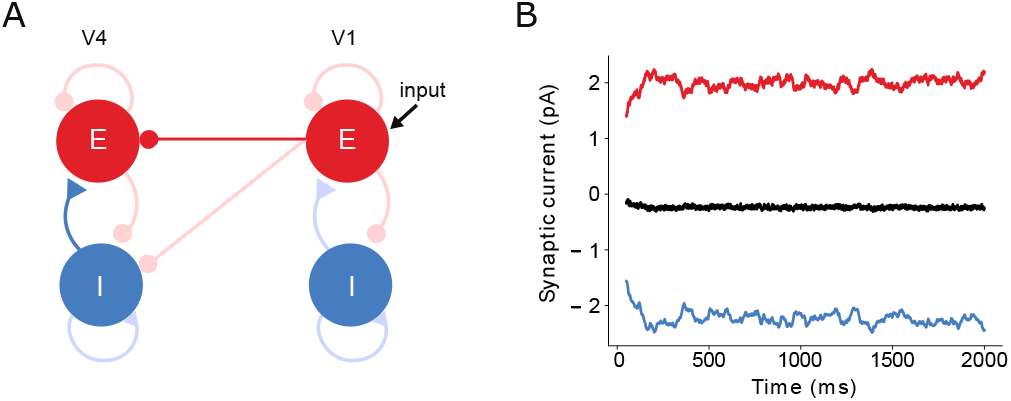
The illustration of detailed balance between inter-areal excitation and intra-areal inhibition. The projection from V1 to V4 is shown as an example. (A), one-way interaction from V1 to V4. V1 receives external Gaussian input. The excitatory population in V4 receives balanced excitatory inter-areal inputs from V1 (dark red) and intra-areal inhibitory inputs from the inhibitory population in V4 (dark blue). Other excitatory and inhibitory interactions in this circuit are colored by light red and blue, respectively. (B), simulation of the synaptic currents received by V4 excitatory population induced by V1 activity. The inter-areal excitatory inputs (red) is balanced with the intra-areal inhibitory inputs (blue), leading to small net inputs (black).

Third, the gradient of *h_i_* parameterizes the gradient of synaptic excitation across areas in the model, supported by the fact that *h_i_* is proportional to the spine count per pyramidal neuron across areas (Chaudhuri et al., 2015; Elston, 2007) in the form of a macroscopic gradient (Wang, 2020). The gradient of synaptic excitation leads to two consequences, (1) it gives rise to the hierarchy of intrinsic timescale for each area while being disconnected to other parts of the cortex, (2) it stabilizes the localization of intrinsic timescale for each area in the presence of long-range connections. From the perturbation analysis and Eq. 11, the degree of eigenvector localization is determined by the competition between the strength of long-range connections encoded in matrix 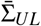 and the spectral gap of matrix 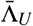. Therefore, the long-range connections tend to delocalize eigenvectors thus break the timescale hierarchy, but the heterogeneity of local recurrent excitation level weakens its effect on eigenvector delocalization in a divisive fashion. In fact, the heterogeneity or randomness in local node properties has been shown to give localized eigenvectors in models of physical system, for instance, a phenomenon known as Anderson localization (Anderson, 1958) that describes the transition from a conducting medium (corresponding to delocalized eigenvectors) to an insulating medium (corresponding to localized eigenvectors). Similar mechanism has been identified in studying the eigenvector localization of a idealized neural network with simple nodes in each cortical area (Chaudhuri et al., 2014).

## Discussion

In this work, we investigated the requirements for the emergence of a hierarchy of temporal response windows in a multi-areal model of the macaque cortex (Chaudhuri et al., 2015). The original model is a nonlinear dynamical system by including a rectified linear f-I curve, and it becomes essentially linear when neural population activities are all above the firing threshold, as happened in our simulations of hierarchical timescale phenomenon. This fact enabled us to define the time constants precisely from the eigenmodes of the connectivity matrix, and carried out a detailed mathematical analysis to identify biologically interpretable conditions. (Rectified) linear models have been broadly used in theoretical and experimental neuroscience studies (Roxin et al., 2005; Binzegger et al., 2009; Jercog et al., 2017). Although microscopic neural activity is nonlinear in general, it has been shown in a recent study (Nozari et al., 2020) that linear models can capture macroscopic cortical dynamics in the resting state more accurately than nonlinear model families, including neural field models for describing the spatiotemporal average of individual neuronal activities. Nonlinear models are more general for capturing neural circuits. However, for a nonlinear model, the time constants of the system are not uniquely defined. A linear model can be understood as a linearization of a nonlinear dynamical system around an internal state such as the resting state of the brain.

In contrast to previous computational models studying the emergence of timescales (Chaudhuri et al., 2014), the model we studied is anatomically more realistic as it incorporates (1) experimental measurements of directed and weighted anatomical connectivity, (2) a gradient of synaptic excitation reflected by spine counts in pyramidal neurons across areas, and (3) both excitatory and inhibitory neuronal populations. By performing rigorous perturbation analysis, we show that the segregation of timescales is attributable to the localization of eigenvectors of the connectivity matrix, and the parameter regime that makes this happen has three crucial properties: (1) a macroscopic gradient of synaptic excitation, (2) distinct electrophysiological properties between excitatory and inhibitory neuronal populations, and (3) a detailed balance between long-range excitatory inputs and local inhibitory inputs for each area-to-area pathway.

The theoretically identified biological conditions for the segregation of timescales enable us to make experimentally testable predictions. First, the condition of the macroscopic gradient of synaptic excitation suggests that shallower gradient of synaptic excitation shall lead to less localized eigenvectors. Consequently, the difference of time constants for a pair of areas shall be larger if their synaptic excitation or hierarchical level are less similar, which can be directly tested in experiments. Second, the condition of distinct electrophysiological properties between excitatory and inhibitory neuronal populations suggests that the change of neuronal physiology will affect the segregation of timescales. This condition can be tested experimentally by using genetic tools to knockdown or knockout specific genes in order to change the firing properties of neurons (Gingras et al., 2014). Third, the condition of detailed balance of excitation and inhibition suggests that areas with unbalanced excitation and inhibition could also alter hierarchical time constants. With growing evidence for E/I imbalance in schizophrenia (Foss-Feig et al., 2017; Jardri et al., 2016), this condition is supported in recent experiments that the intrinsic time constants of schizophrenia patients have been substantially changed (Wengler et al., 2020). And this condition can be further tested in animal models with genetic tools to disturb the excitation-inhibition balance (Gatto and Broadie, 2010).

Beyond neuroscience, eigenmode localization is an important phenomenon in nature that has aroused broad interests. A prominent example is known as Anderson localization in physics (Anderson, 1958), finding that the eigenmodes of the Schrödinger equation that govern the wave function of electrons can be spatially localized with heterogeneous potentials. Anderson localization successfully explains the transition from metal to insulator given impurities or defects in materials. In addition to that, light (Wiersma et al., 1997), acoustic waves (Hu et al., 2008), seismic waves (Larose et al., 2004) and others (Billy et al., 2008; Choi et al., 2017; White et al., 2020) in heterogeneous medias have also been found to exhibit localized modes. The interactions in these systems are short-range, in contrast to brain circuits with an abundance of long-range connections. Here we found that eigenmodes can also be localized in the cortical network with similar but different mechanisms from the aforementioned systems. The localization in the cortex also requires heterogeneous local properties (i.e., recurrent excitation level), and it requires additional distinct properties of the excitatory and inhibitory neurons and their balanced interactions. Our work thus provided another example of eigenmode localization in natural systems and demonstrated that the intrinsic properties of “particles” in the system could be important to eigenmode localization.

It is worth mentioning that, although the specific pattern of inter-areal connectivity does not affect the eigenvector localization substantially based on the perturbation analysis, it shapes the timescale hierarchy qualitatively. In particular, the timescale hierarchy does not exactly follow monotonically the areal anatomical hierarchy in the presence of long-range connections, as shown in Fig. 1C. Furthermore, within a brain region time constants are heterogeneous across individual neurons (Bernacchia et al., 2011; Cavanagh et al., 2020). To better relate the model with experimentally observed timescales in various specific cortical areas, the roles of long-range connections, cell types and other circuit properties require further elucidation.

It has been noticed that the neuronal activity propagates along the hierarchy with significant attenuation in the model of Ref. (Chaudhuri et al., 2015). The attenuation can be alleviated by tuning the model parameters to the regime of strong global balanced amplification (GBA) (Joglekar et al., 2018) (see parameters in Sec. *Materials and Methods*). Balanced amplification was originally introduced for a local neural network, associated with strong non-normality of the system where eigenmodes are far from being orthogonal with each other (Murphy and Miller, 2009). A quantity called *κ* measures the degree of non-normality of a matrix (Trefethen and Embree, 2020) (*κ* = 1 for a normal matrix; the larger the *κ* value, the more non-normal the system). We have *κ* = 4.35 for the original model (Chaudhuri et al., 2015), which is thus only slightly non-normal. By contrast, *κ* = 96.58 for the model in the strong GBA regime. Therefore, the enhancement of signal propagation in the model correlates with the increase of the non-orthogonality of the eigenvectors, or the non-normality of the connectivity matrix. In the strong GBA regime, *δ* ≈ 0.38, which is ten times larger than its original value, suggesting that the detailed balance condition is less well satisfied. In such a case, the localization of time scales may no longer exist in this linear model. The situation is different in nonlinear models (Chaudhuri et al., 2015; Mejias and Wang, 2020; Chien and Honey, 2020), where inputs may be amplified by strongly recurrent circuit dynamics to enhance signal propagation or routing of information is selectively gated (for a subset of connection pathways in a goal-directed manner) (Vogels and Abbott, 2009; Wang and Yang, 2018), while the conditions for a timescale hierarchy are satisfied. For a nonlinear system, however, eigenmodes can be defined only with respect to a particular network state. Consequently, the time constants observed in single neurons are no longer unique and may differ, for instance, when the brain is at rest or during a cognitive process. It remains to be seen to what extent the conditions identified here hold in various brain’s internal states, while the precise pattern of timescales can be flexibly varied to meet behavioral demands.

## Acknowledgements

We thank Y. Xiao and H. Wang for helpful discussions. This work was supported by National Key R&D Program of China 2019YFA0709503, Shanghai Municipal Science and Technology Major Project 2021SHZDZX0102, National Natural Science Foundation of China Grant 11901388, and Shanghai Sailing Program 19YF1421400 (S.L.), James Simons foundation grant 543057SPI (X.-J.W.)

## Author Contributions

S.L. and X.-J.W. designed the research, S.L. performed the work, S.L. and X.-J.W. wrote the paper.

## Materials and Methods

### Model parameters

In the macaque cortical network model, we set *τ_E_* = 20 ms, *τ_I_* = 10 ms, *β_E_* = 0.066 Hz/pA, *β_I_* = 0.351 Hz/pA, *w_EE_* = 24.4 pA/Hz, *w_IE_* = 12.2 pA/Hz, *w_EI_* = 19.7 pA/Hz, *w_II_* = 12.5 pA/Hz, *μ_EE_* = 33.7 pA/Hz, *μ_IE_* = 25.5 pA/Hz and *η* = 0.68. We set *w_EI_* = 25.2 pA/Hz and *μ_EE_* = 51.5 pA/Hz for the strong balanced amplification regime (Joglekar et al., 2018) introduced in Sec. Discussion. Some of the parameters are derived from experimental measurements of primary visual cortex (Binzegger et al., 2009). The FLN values are obtained from the experimental measurements of macaque cortical connectivity (Markov et al., 2014a). The hierarchy values *h_i_* of each cortical area are obtained by fitting a generalized linear model that assigns hierarchical values to areas (Chaudhuri et al., 2015) such that the differences in hierarchical values predict the supragranular layer neurons (SLNs) measured in experiment (Markov et al., 2014b).

## Code Availability

Upon publication, the python code for model simulation and perturbation analysis will be made publicly available on Github (https://github.com/songting858).

## Appendix

### Proposition 4

*Given*

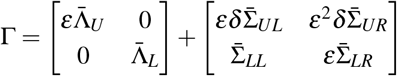

*and*

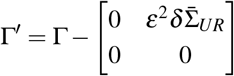

*with* 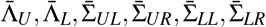 *being* 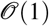, *and A_U_ has n simple eigenvalues. If λ and **X** are the eigenvalue and eigenvector of* Γ *respectively, then there exists λ′ and **X**′ being the eigenvalue and eigenvector of* Γ′ *respectively, such that* 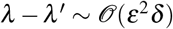 *or higher order*, 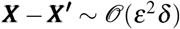 *or higher order*.

**Proof:** The eigenvalue of Γ denoted by *λ* can be solved as the root of the characteristic polynomial

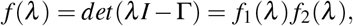

where

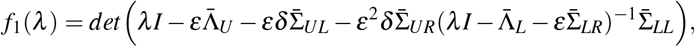

and

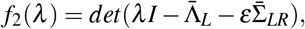

here *det*(·) is the determinant of a matrix, and *I* is the identity matrix.

For the eigenvalues that solve *f*_1_(*λ*) = 0, if we define 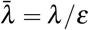, then 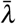 solves

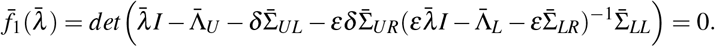

If we set *β* = *εδ*, then 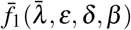 is analytic with respect to 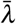, *ε*, *δ*, *β*. In addition, as 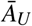 has *n* different eigenvalues denoted by 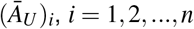, we have 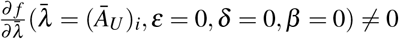 for *i* = 1,2, …, *n*. According to the implicit function theorem, 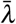 is analytic near 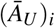 with respect to *ε*, *δ*, *β*. Therefore, the eigenvalue of Γ can be expanded as 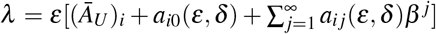 with coefficients *a_ij_*(*ε, δ*) being analytic with respect to *ε* and *δ*. And the eigenvalue of Γ′ denoted by *λ*′ can thus be obtained by setting *β* = 0 as 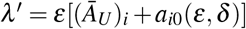. Consequently, we have 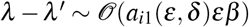 which is of order 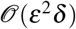 or higher. For the eigenvalues that solve *f*_2_(*λ*) = 0, they do not depend on *β* as *f*_2_(*λ*) does not include *β*. Consequently, these *n* eigenvalues of Γ and Γ′ are identical.

For eigenvector *X* of matrix Γ, if we set *γ* = *ε*^2^*δ*, then Γ is a analytic function of *ε*, *δ*, and *γ*. Accordingly, its eigenvector *X* is analytic with respect to *ε*, *δ*, *γ* (Kato, 1966). In particular, we can expand 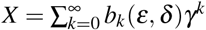 with coefficient *b*(*ε, δ*) being analytic. And the eigenvector *X*′ of matrix Γ′ can thus be obtained by setting *γ* = 0 as *X*′ = *b*_0_(*ε, δ*). Therefore, the difference is 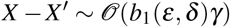 which is of order 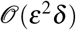 or higher.

## Notes

### Competing Interest Statement

The authors have declared no competing interest.

